# Potential of *in vivo*- and *in vitro*-cultured entomopathogenic nematodes to infect *Lobesia vanillana* (Lepidoptera: Tortricidae) under laboratory conditions

**DOI:** 10.1101/2020.11.09.374066

**Authors:** Francois du Preez, Antoinette Paula Malan, Pia Addison

**Author notes:** Corresponding author, (FDP).

## Abstract

The *in vivo*- and *in vitro*-cultured South African entomopathogenic nematodes (EPNs), *Steinernema yirgalemense* and *Steinernema jeffreyense* (Rhabditida: Steinernematidae), were evaluated against larvae and pupae of *Lobesia vanillana* in laboratory bioassays. For larvae, high mortality was observed for all treatments: *In vitro*-cultured *S. yirgalemense* (98%) performed better than *S. jeffreyense* (73%), while within *in vivo* cultures, there was no difference between nematode species (both 83%). No significant difference was detected between *in vivo*- and *in vitro* cultures of the same nematode species. The LD_50_ of the *in vitro*-cultured *S. yirgalemense*, the best performing species, was 7.33 nematodes per larva. Mortality by infection was established by dissecting cadavers and confirming the presence of nematodes, which was > 90% for all treatments. Within *in vitro* cultures, both *S. yirgalemense* and *S. jeffreyense* were able to produce a new cohort of infective juveniles from *L. vanillana* larvae. Pupae, however, were found to be considerably less susceptible to EPN infection. The relative success of local *in vivo*- and *in vitro*-cultured EPN species against a tortricid species in laboratory assays is encouraging for further research and development of this technology.

## Introduction

*Lobesia vanillana* (De Joannis) (Lepidoptera: Tortricidae) occurs throughout the Afrotropical region [1] and appears to be polyphagous. Apart from limited information relating to taxonomy and locality, very few records exist in literature. It has been reported as a pest of *Mangifera* sp. (Anacardiaceae) and *Vanilla planifolia* (Orchidaceae) [2], from which it gets its name [3].

Natural enemies for *L. vanillana* are unknown. Biological control agents, including *Bacillus thuringiensis* (Bt) (Bacillalus: Bacillaciae) and mating disruption products, have been registered and used in Europe on *Lobesia* species [4]. Natural enemies such as *Trichogramma* parasitoid wasps (Hymenoptera: Trichogrammatidae) have been identified in Europe [5].

Diapausing pupae of *Lobesia botrana* overwinters under the bark of vines and emerge the following spring [6], and *L. vanillana* is suspected to follow a similar life cycle. Observations by the author suggests that larvae and pupae of *L. vanillana* do not have a soil-dependent life stage.

Insect parasitic nematodes, or entomopathogenic nematodes (EPNs), occur in soils across the world and are natural enemies of many insect species. EPNs of the family Steinernematidae (Rhabditida) are associated with the symbiotic bacteria of *Xenorhabdus* (Enterobacteriales: Enterobacteriaceae) [7]. Infective juveniles (IJs) of these species release their symbiotic bacteria shortly after penetrating the haemocoel of their target insect, causing mortality within 48 h [8], depending on the number of nematodes that penetrated, and the size of the insect host [9]. The virulence of these pathogens is usually correlated with host abundance and the associated microclimate [10]. In agricultural applications, they can be used as inundative control (similar to chemical insecticides), but in favourable conditions, they can successfully establish to provide persistence [9].

*Steinernema* has been previously evaluated against lepidopteran pests, under both laboratory and field conditions. Relevant EPN biocontrol research in South Africa includes that of the above-ground diapausing larval population of the codling moth, *Cydia pomonella* L. (Lepidoptera: Tortricidae) [11–14] and the soil stages of false codling moth *Thaumatotibia leucotreta* (Meyrick) (Lepidoptera: Tortricidae) [15–18].

*Steinernema yirgalemense* Nguyen, Tesfamariam, Gozel, Gaugler & Adams and *Steinernema jeffreyense* Malan, Knoetze & Tiedt have been successfully mass-produced using *in vitro* liquid culture methods [19,20]. Promising results were obtained against false codling moth, using both *in vivo*- and *in vitro*-cultured nematodes, with no significant difference between culture types in laboratory and field trials [18]. The *in vitro* liquid mass production of EPNs is much more cost and labour effective than *in vivo* production, the latter of which is better suited for small-scale experiments and insecticidal applications [21].

Insects without a soil stage may be more susceptible to EPNs, as they may not have had the opportunity to evolve the resistance necessary to protect themselves from nematode infections. This weakness of above-ground pest defence mechanisms against microbiological pathogens can thus be exploited to provide biological control, for example, as with previous research on mealybugs, and the addition of adjuvants to nematode suspensions [18,22–26] has shown.

The aim of this study was to evaluate the pathogenicity of *in vivo*- and *in vitro*-cultured *S. yirgalemense* and *S. jeffreyense* against the larvae and pupae of *L. vanillana*. Screening was conducted under optimum laboratory conditions and mortality by infection confirmed. The LD_50_ of the most efficient species was determined. Additionally, reproduction of the nematode in the insect cadaver was investigated.

## Materials and methods

### Source of insects

Larvae were obtained from a mass rearing facility in Stellenbosch. They were fed an agar-based modified codling moth diet [27] and kept at a constant 25°C with a 18:6 h light-dark cycle. Adults were placed next to a window to receive natural indirect sunlight, and were given a cotton ball, dipped in a 2% sugar-water solution, for nourishment.

*Tenebrio molitor* L. (Coleoptera: Tenebrionidae) larvae, used for the *in vivo* culture of EPNs, were cultured in the laboratory on bran and carrots, in vented containers according to the technique of Van Zyl and Malan (2015) [28].

### Source of nematodes

*In vitro*-cultured *S. yirgalemense* (157-C) (GenBank accession number EU625295) [15,19] and *S. jeffreyense* (J194) (KC897093) [29] were cultured according to the technique of Dunn *et al.* [20]. *In vivo* cultured nematodes, of the same two species, were cultured *in vivo* using *T. molitor* larvae. Approximately 20 ml of each nematode species were prepared, counted and placed in culture flasks. Flasks were stored horizontally, shaken biweekly to discourage nematode clumping and to aerate the suspension, and were used within two months. The viability of cultures was evaluated by inspecting nematode samples for motility and mortality, using a standard stereo microscope.

### Bioassay protocol

The test arena consisted of 24-well bioassay trays (CellStar TC, Cat No 662160, Greiner Bio-One, Frickenhausen, Germany). Filter paper discs (12.7 mm diam., Grade FN 30, Cat No 3.526.012-7, Ahlstrom-Munksjö, Bärenstein, Germany) were added to each alternate well and alternate row, on which IJs were inoculated at a predetermined concentration [30], by pipetting 50 μl of the nematode suspension onto the filter paper. Each treatment consisted of four 24-well bioassay plates, with six *L. vanillana* individuals per tray, for a total of 24 larvae per treatment. For larval bioassays, glass rectangles were placed between the lid and the tray to prevent the escape of larvae during the incubation period.

Trays were stacked in closed 2-L plastic ice cream containers, each with moistened paper towels at the bottom, to provide an environment of high humidity, then incubated at 25°C for 48 h in the dark. Mortality by EPN infection was assessed by gently prodding the insect with a dissection needle and evaluating the integrity and colour of cadavers.

Larvae were carefully rinsed using a handheld water jet to rid the cadaver of surface nematodes. Cadavers were then either placed on a modified White Trap [31] to evaluate *in vivo* nematode production, or dissected to evaluate nematode infection.

### Susceptibility of larvae

Following the described protocol, bioassays consisted of two treatment groups (*in vivo* and *in vitro*), inoculated with *S. yirgalemense* and *S. jeffreyense*, and one control group treated with water only. Larvae were inoculated with 100 IJs/50 μl for all treatments. Controls received 50 μl distilled water only. The experiment was repeated on a different date, with a different batch of nematodes.

### Susceptibility of pupae

Screening for the susceptibility of *L. vanillana* pupae was according to the described protocol, for the two treatment groups (*in vivo* and *in vitro*), inoculated with *S. yirgalemense* and *S. jeffreyense*, and one control group treated with water only. Pupae were inoculated with 200 IJs/50 μl for all treatments. Controls received 50 μl distilled water only. The experiment was repeated on a different date with a different batch of nematodes.

### Dose-response for larvae

Following the results of susceptibility bioassays for *L. vanillana* larvae, the most effective EPN species was selected for calculating LD_50_ estimates. The same inoculation procedure as described in the bioassay protocol was followed. Logarithmic dosages translated to 100, 50, 25, 12.5, 6.25 and 0 IJs/larva, respectively.

### Nematode penetration and reproduction

To assess the infectivity of nematodes, half of the infected *L. vanillana* larvae from the susceptibility bioassay were incubated at 25°C, at high humidity, and dissected 18-36 h after the initial 48 h incubation period. This allowed nematodes to grow within the cadaver, which improved their visibility for counting. Infected larvae were placed singly on a petri dish, with a droplet of distilled water to suspend the cadaver contents, and dissected with the aid of a Leica MZ75 stereo microscope. The presence and number of nematodes within the cadaver was recorded.

To assess the reproductive ability of nematodes within the infected *L. vanillana* larvae, the remaining cadavers of the susceptibility bioassay were placed in 90 mm diam. plastic petri dishes with one filter paper circle, (85 mm, Grade 1 Whatman, CAT no. 1001-085, GE Healthcare Life Sciences) moistened with 800 μl of distilled water. Approximately 10-12 cadavers were placed per petri dish. With the top removed, plastic petri dishes were transferred to 150 mm glass petri dishes, the bottom of which contained just enough distilled water to not float the plastic petri dishes, and covered with the glass lid. The resulting IJ suspension was harvested three times during the course of 45 days, and the concentration determined as nematodes per volume, divided by the number of larvae on the trap, to give the number of nematodes produced per larva. Distilled water was added to the glass- and plastic petri dishes as needed.

### Data analysis

Data were analysed in Microsoft Excel 2016 for descriptive statistics and processed in Statistica 13.3 [32] for comparative analysis. Probit analysis and LD_50_ estimates were calculated using NCSS [33]. For larval susceptibility bioassays, residuals of the mortality response were considered normally distributed (Shapiro-Wilk’s W = 0.967, p = 0.187), permitting the use of a factorial ANOVA and Fisher’s LSD post-hoc test to evaluate responses between nematode production types and between nematode species. For pupal susceptibility bioassays, residuals of the mortality response were considered normally distributed, despite having a significant p-value (Shapiro-Wilk’s W = 0.923, p = 0.004), by examining relevant normality graphs. All analyses were evaluated for, and passed, Levene’s test for homogeneity of variances (p > 0.05). Results are given as the mean response for all repetitions ± standard error, unless otherwise specified.

## Results

### Susceptibility of larvae

Mortality of *L. vanillana* larvae significantly interacted with EPN species (F = 105.167, p < 0.005), but not with EPN culture type (F = 0.0848, p = 0.772). There was no significant interaction between EPN species and EPN culture type (F = 2.888, p = 0.0668).

Within *in vitro* cultures, *S. yirgalemense* (97.88% ± 2.13%) was significantly (p < 0.01) more effective against *L. vanillana* larvae than *S. jeffreyense* (72.88% ± 6.21%), while both were significantly (p < 0.01) more effective than the control (8.38% ± 6.31%). Within *in vivo* cultures, *S. yirgalemense* (83.25% ± 6.28%) and *S. jeffreyense* (83.38% ± 5.4%) were significantly different from the control (16.63% ± 7.02%), but not from each other (Fig 1).

**Fig 1.**
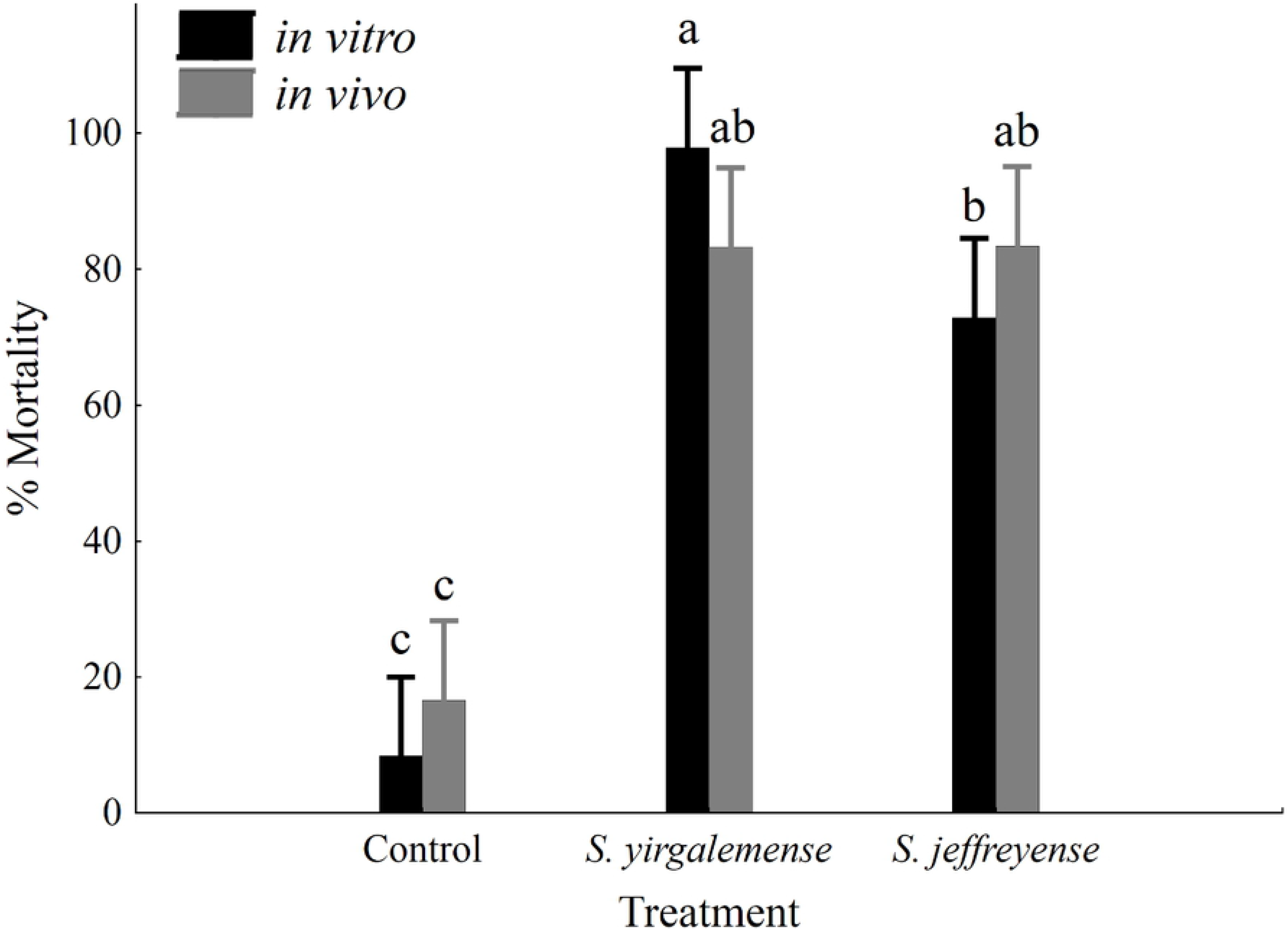
Mortality of *Lobesia vanillana* larvae 48 h after inoculation with nematodes. Percentage insect mortality (95% confidence intervals) of *L. vanillana* larvae, 48 h after inoculation with *in vitro*- and *in vivo*-cultured infective juveniles of *Steinernema yirgalemense* and *S. jeffreyense*, at a concentration of 100 IJs/50 μl. Vertical bars were calculated using least square means. Different letters on the bars denote statistical significance, calculated using Fisher’s LSD (MSE = 267.64; df = 42; p < 0.05).

In the case of *L. vanillana* pupae, mortality significantly interacted with EPN species (F = 4.798, p = 0.0133), but not with EPN culture type (F = 0.0787, p = 0.781). There was no significant interaction between EPN species and EPN culture type (F = 2.438, p = 0.0996).

*In vitro-*cultured *S. yirgalemense* performed significantly better (14.58% ± 3.78%) than its control (0.00% ± 0.00%). *In vivo*-cultured *S. yirgalemense* (6.25% ± 3.05%) performed slightly better than its control (2.08% ± 2.08%), as did *in vitro*- and *in vivo*-cultured *S. jeffreyense* (4.17% ± 2.73% and 8.33% ± 4.45%, respectively), but none of these differences were statistically significant (Fig 2).

**Fig 2.**
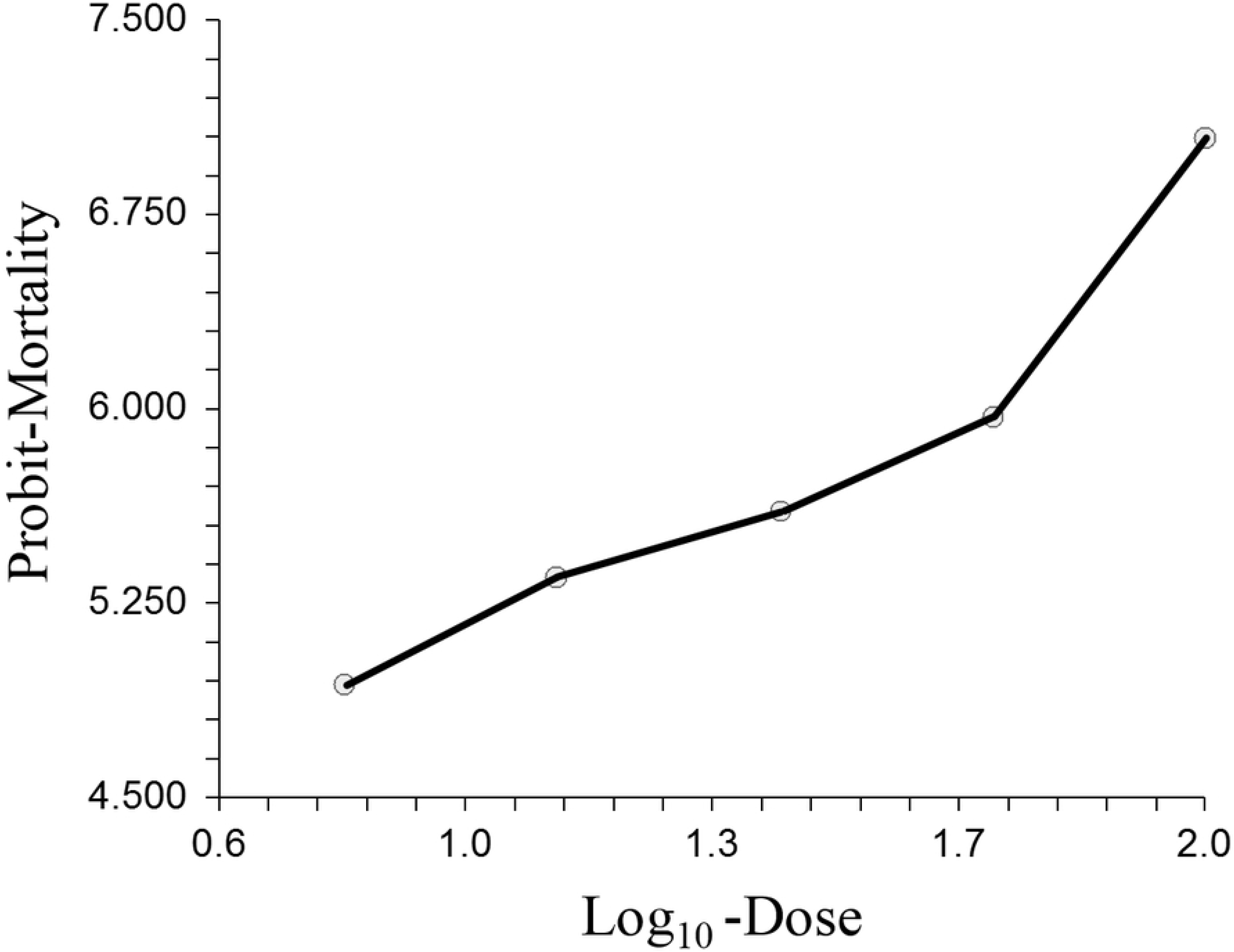
Mortality of *Lobesia vanillana* pupae, 48 h after inoculation with nematodes. Percentage mortality (95% confidence intervals) of *L vanillana* pupae, 48 h after inoculation with *in vitro*- and *in vivo*-cultured infective juveniles (IJ) of *Steinernema yirgalemense* and *S. jeffreyense*, at a concentration of 200 IJs/50 μl. Vertical bars were calculated using least square means. Different letters on the bars denote statistical significance, calculated using Games-Howell (MSE = .00736; df = 42; p < 0.05).

Low EPN performance in susceptibility bioassays against *L. vanillana* pupae resulted in an insufficient number of cadavers for dose-response, penetration and reproduction analyses.

### Dose-response for larvae

The data fits the Probit model well (Chi-Square = 1.29; DF = 3; Prob. level = 0.73). Lethal dosage estimates for *in vitro*-cultured *S. yirgalemense* were calculated as follows: LD_25_ = 2.37 ± 1.37; LD_50_ = 7.335 ± 2.485; LD_90_ = 62.761 ± 23.224 and LD_95_ = 115.339 ± 57.426 nematodes per larvae. The Probit model may be expressed by the linear function *P* = 3.81 ± 0.466 + 1.375*x* ± 0.344, where *P* is Probit-Mortality and *x* is Log10-Dose (Fig 3).

**Fig 3.**
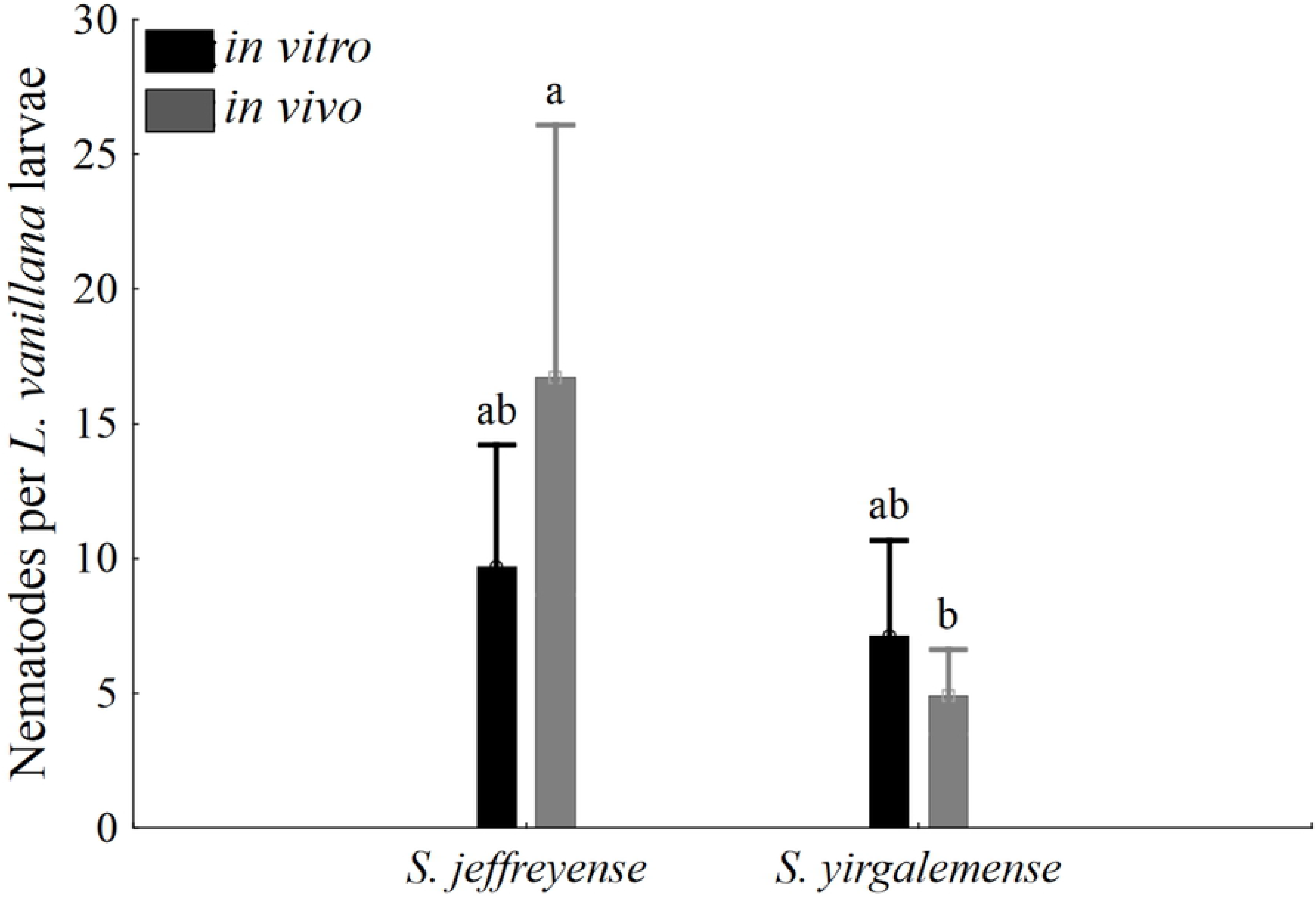
Mortality of *Lobesia vanillana* 48 h after inoculation with different dosages of nematodes. Probit mortality of *L. vanillana* larvae 48 h after inoculation at different dosages (100, 50, 25, 12, 6 and 0 IJs/larva) of *in vitro*-cultured *Steinernema yirgalemense*. The LD_50_ was estimated as 7.335 ± 2.485 nematodes.

### Nematode penetration and reproduction

In all treatments, > 90 % of the cadavers had nematodes present (Fig 4). A higher percentage of nematodes penetrated *L. vanillana* larvae in the *in vivo*-cultured treatments versus the *in vitro* counterparts, however this difference was not significant (p = 0.978).

**Fig 4.**
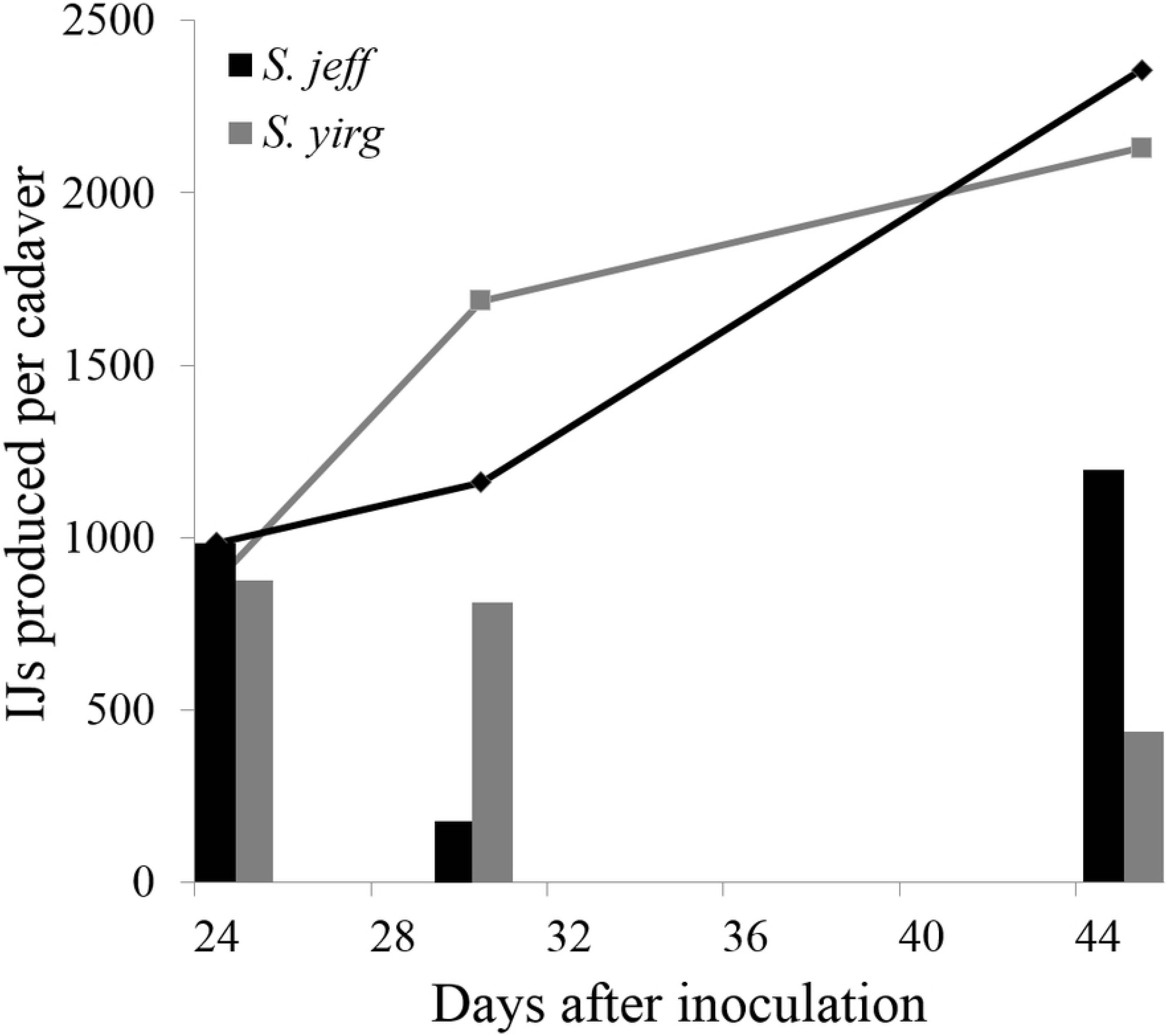
*Lobesia vanillana* cadavers infected with nematodes. Percentage of *L. vanillana* cadavers with nematodes present following susceptibility bioassays of *in vitro*- and *in vivo*-cultured *Steinernema yirgalemense* (*S. yirg*) and *S. jeffreyense* (*S. jeff*). No significant differences were found between treatments.

Chi-square analysis revealed no significant differences between IJ penetration within cadavers and nematode culture type or nematode species. Residuals of nematode counts failed the normality assumption (Shapiro-Wilk’s W = 0.771, p < 0.01), but relatively large sample sizes per group (n ≥ 22) allowed for an ANOVA bootstrap analysis. There was a significant difference in the average number of nematodes per cadaver within the *in vivo* culture type, between *S. jeffreyense* (17.091 ± 0.187) and *S. yirgalemense* (4.917 ± 0.334) (Bootstrap p = 0.045), while within the *in vitro* culture type, there was no significant difference between them (Fig 5).

**Fig 5.**
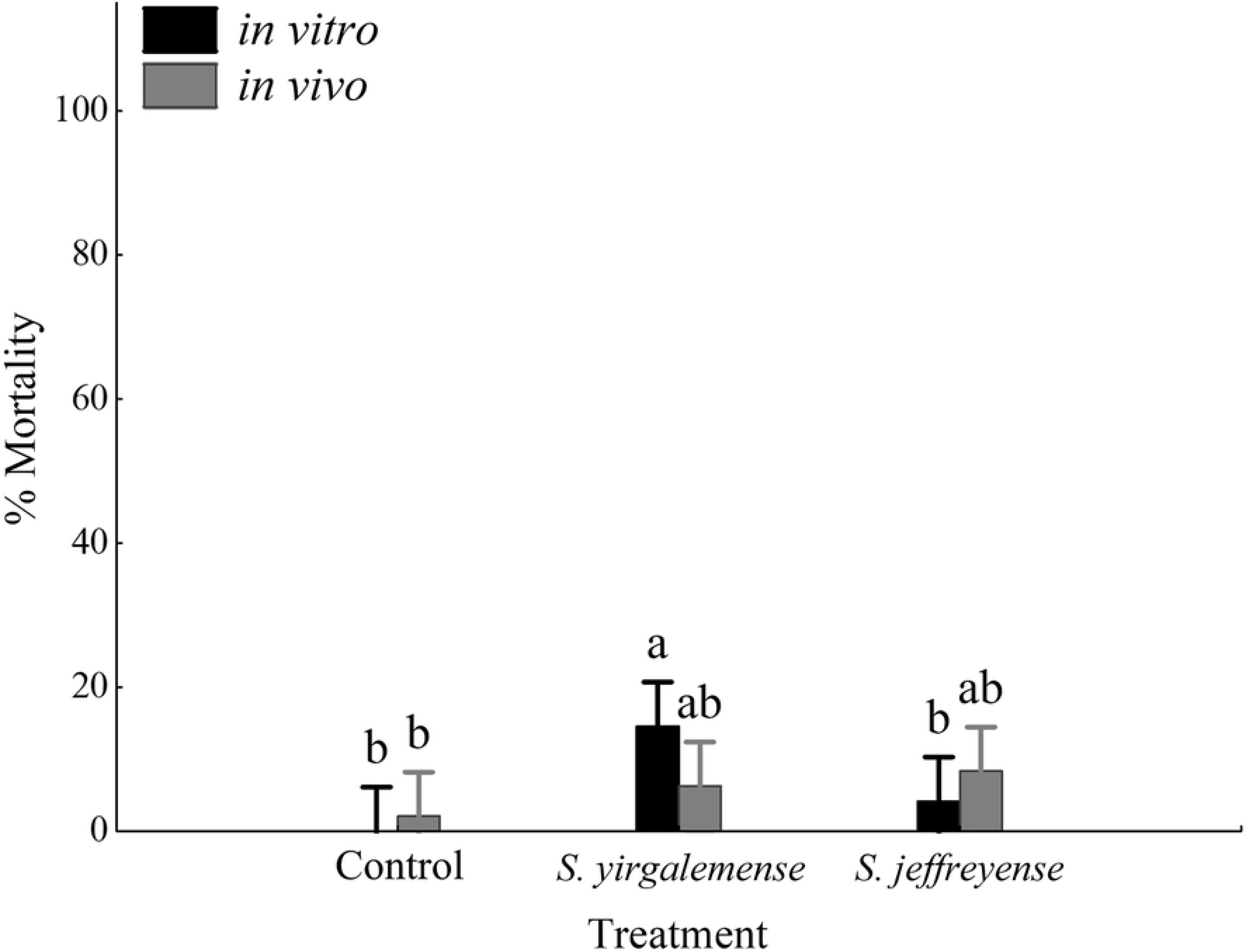
Number of nematodes penetrated *Lobesia vanillana* larvae. Average number of nematodes penetrated per *L. vanillana* larvae, following susceptibility bioassays, using *in vitro*- and *in vivo*-cultured *Steinernema yirgalemense* and *S. jeffreyense*. Different letters on the bars denote statistical significance (p < 0.05), calculated using Bootstrap.

Both *in vitro*-cultured *S. yirgalemense* and *S. jeffreyense* nematodes were able to produce IJs from cadavers following the susceptibility bioassay. Cumulative production after 45 days totalled 2 130 IJs for *S. yirgalemense* and 2 356 IJs for *S. jeffreyense*, per insect cadaver (Fig 6).

**Fig 6.**
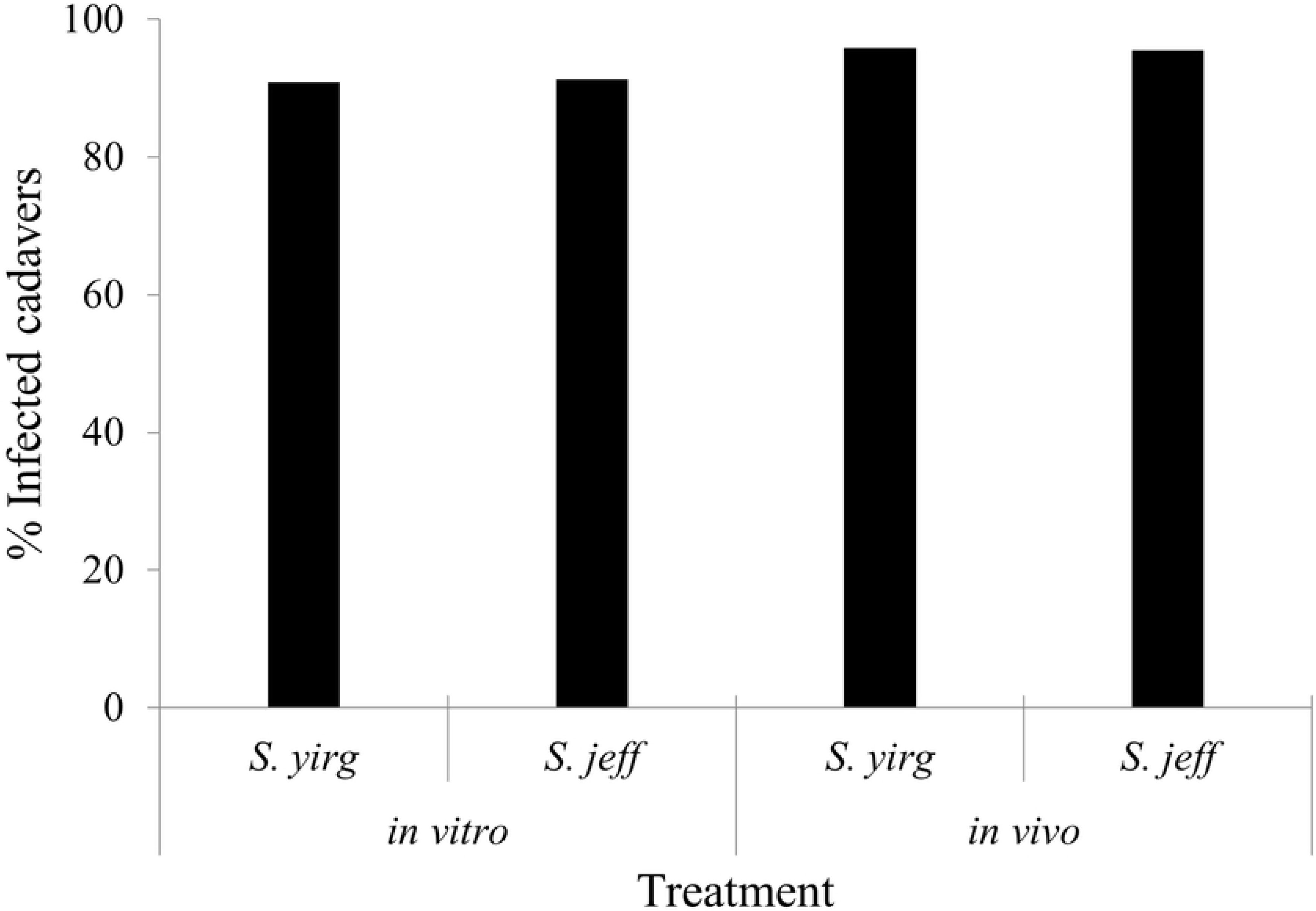
Number of nematodes produced per *Lobesia vanillana* larvae. Average number of infective juveniles (IJs) produced per *L. vanillana* larvae, following susceptibility bioassays, using *in vitro*-cultured *Steinernema yirgalemense* (*S. yirg*) and *S. jeffreyense* (*S. jeff*). Lines indicate cumulative production.

## Discussion

This was the first study evaluating EPNs for the control of *L. vanillana*. For larvae, all treatments resulted in exceptionally high mortality. Specifically, within *in vitro* cultures, *S. yirgalemense* performed significantly better than *S. jeffreyense*, while within *in vivo* cultures, there were no significant difference in mortality between species. There was no significant difference found between *in vitro* and *in vivo* cultures of the same species.

Low mortality was observed for all treatments against pupae. Only *in vitro*-cultured *S. yirgalemense* performed significantly better (15%) than the control, while the other treatments had lower mortality (< 9%), not significantly different from each other, nor from the control. In general, lepidopteran pupae are less susceptible to infection, as indicated by Malan *et al.* [15] for false codling moth. However, Steyn *et al*. [18] suggested that the age of pupae have an effect on susceptibility, with older fully formed, hardened pupae, being less susceptible.

Observations found *Lobesia vanillana* to be relatively small (adults ± 6 mm in length), compared to other tortricid moths previously evaluated against EPNs, such as codling moth and false codling moth [34]. The LD_50_ of *in vitro*-cultured *S. yirgalemense* against *L. vanillana* larvae was estimated as 7 IJs/larva.

In a study by De Waal *et al.* [11], both *in vivo*-cultured *S. yirgalemense* and *S. jeffreyense* were evaluated against the above-ground diapausing codling moth larvae, at half the concentration used for *L. vanillana* in the present study (50 IJs/larva). Both nematode species resulted in a mortality of close to 100%. However, Odendaal *et al.* [14] found that in a semi-field spray trial, *S. jeffreyense* performed better than *S. yirgalemense*, contrary to what was expected.

Mortality, caused by nematode infection, was confirmed by dissecting *L. vanillana* larvae and evaluating the presence and number of nematodes. Within *in vivo* cultures, there was a significant difference between *S. jeffreyense* (17 nematodes per cadaver) and *S. yirgalemense* (five nematodes per cadaver), but no significant difference between the species of *in vitro* cultures. It was expected that the number of IJ penetrated would be higher in the case of *S. yirgalemense*, as it is a smaller IJ (± 635 μm) [35] when compared to the body length of *S. jeffreyense* (> 900 μm) [29].

Visual observation of larval cadavers directly after susceptibility bioassays, revealed that few of the final instar insect larvae treated with *S. yirgalemense* managed to pupate in their trays, with little webbing present, compared to those treated with *S. jeffreyense*, suggesting that *S. yirgalemense* is faster-acting than *S. jeffreyense*, in laboratory bioassays at least, but more research is needed to support this theory.

The *in vitro* cultures of both *S. yirgalemense* and *S. jeffreyense* had the ability to produce a new cohort of IJs, and after 45 days produced an estimated 2 130 IJ and 2 356 IJs per cadaver, respectively. Generally, the larval stages of lepidopterans were found to support nematode infection and reproduction [34], and more so when insects are believed to not have a soil stage. In rare cases, especially where insects have soil stages, a type of resistance against nematodes can develop, such as the case with woolly apple aphid, *Eriosoma lanigerum* (Hausmann) (Hemiptera: Aphididae) [36].

Local research established *in vitro* liquid culture methods for *H. zealandica*, *S. jeffreyense* and *S. yirgalemense* [19,20,37] while research on the formulation, packaging and storage of these species is still ongoing [38–40]. *In vivo*-cultured IJs can provide affordable, high quality nematodes that are easy to culture, but only on a small scale [21]. Increased complexity, risk, labour and running costs are prohibitive when scaling towards mass-production [41]. The start-up capital and complexity of *in vitro* production methods are excessive for small-scale use, but for mass-production and augmentative releases where a large number of nematodes is required, it is the most cost-effective solution [21]. Results from this study indicate that the quality of *in vitro*-cultured nematodes are comparable to those cultured *in vivo*, the latter of which is considered the “more natural” method. Previous studies by Ferreira *et al.* [19,37] evaluated the efficacy of *in vitro*- and *in vivo*-cultured *S. yirgalemense* and *H. zealandica* against the greater wax moth, *Galleria mellonella* (L.) (Lepidoptera: Pyralidae), and found *in vivo*-cultured nematodes to cause significantly higher mortality than their *in vitro*-cultured equivalents.

Steyn *et al.* [18] evaluated *S. jeffreyense* and *S. yirgalemense*, both *in vivo* and *in vitro*, cultured under the same conditions as the present study, and assessed their mortality against false codling moth, both in the laboratory and in the field. Using half the concentration of IJs used in the current study (50 IJs/larva), high mortality of false codling moth was found in the laboratory, compared to semi-field applications which resulted in a mortality of ≈ 70%. Similar to the present study, no significant difference was found between *in vivo*- and *in vitro*-cultured nematodes of these two species [18].

The isolate of *S. yirgalemense* demonstrated high efficacy against other key insect pests of various fruit crops in South Africa, including banded fruit weevil [42], fruit fly [43] and mealybug [24], and has been prioritised for commercialisation [39,40].

Environmental parameters and fluctuations (temperature, relative humidity, wind, UV, etc.) may affect the interactions between the nematode and its insect host, as well as their survival, dispersal and fecundity. Lower efficacies can be expected in field trials, due to harsher conditions and environmental incompatibilities relative to the laboratory bioassay protocol. For example, in codling moth trials, lower temperature resulted in slower and lower mortalities compared with warmer temperatures [14]. The origin of nematode cultures may also influence their performance, due to their ability (or inability) to adapt to different climates. For example, following results from field trials, Odendaal *et al.* [13] proposed that because *S. yirgalemense* originated from Mpumalanga [15], it may not be well adapted to Mediterranean climates.

Larvae of *L. vanillana* in the present study were sourced from a laboratory colony established in February 2018 from field-collected individuals, and the susceptibility of these individuals to EPNs may differ slightly to that of the natural field population. More nematode species, especially *H. zealandica* and other native species that show effective control against lepidopteran pests, can be evaluated in future research to establish a nematode susceptibly profile for *L. vanillana*. In addition, the application of nematode formulations to the canopy and soil of orchards may have the ability to control multiple pests simultaneously [22,25,26,44–46], especially when used in an integrated pest management programme.

Results of the present study thus indicate that *in vitro*- and *in vivo*-cultured *S. yirgalemense* and *S. jeffreyense* nematodes are able to infect and kill the larvae of *L. vanillana*, and that these *in vitro* cultures are able reproduce within this host and produce a new cohort of IJs, capable of finding and infecting new hosts. The relative success of *S. yirgalemense*, *S. jeffreyense* and other local EPN species against South African tortricid species in laboratory and field assays, the ability to produce nematodes using *in vitro* liquid culture techniques, and the industry demand for such products, encourages further research and development of this technology.

## Acknowledgements

The authors wish to thank InsectUS for assistance in the rearing of insects.

## Author contributions

**Conceptualization**: Francois du Preez, Antoinette Malan, Pia Addison

**Formal Analysis**: Francois du Preez

**Funding Acquisition**: Pia Addison

**Investigation**: Francois du Preez

**Methodology**: Antoinette Malan, Francois Du Preez

**Project Administration**: Pia Addison

**Supervision**: Pia Addison, Antoinette Malan

**Writing – Original Draft Preparation**: Francois Du Preez

**Writing – Review & Editing**: Francois du Preez, Antoinette Malan, Pia Addison

